# An Optimized LIVE/DEAD Assay Using Flow Cytometry to Quantify Post-Stress and Antifungal-Treatment Survival in Diverse Yeasts

**DOI:** 10.1101/2025.04.14.648826

**Authors:** Hanxi Tang, Jinye Liang, Bin Z. He

## Abstract

Quantifying post-stress survival in yeasts is crucial for biological, biomedical, and industrial research. Traditional methods like Colony Forming Unit (CFU) assays are labor-intensive and time-consuming. In this study, we systematically characterize a two-dye (SYTO 9 / Propidium Iodide) LIVE/DEAD assay coupled with flow cytometry to rapidly and scalably quantify post-stress survival in diverse yeast species. By optimizing staining buffer, dye concentrations and staining time, we minimized artifacts and improved reproducibility. Notably, we identify an “Intermediate” population containing damaged cells with enhanced SYTO 9 uptake but little propidium iodide (PI) accumulation under sublethal stress, providing finer gradations of cellular damage compared to CFU or PI staining alone. We demonstrate the assay’s applicability across *Candida glabrata*, *Saccharomyces cerevisiae*, and *Candida albicans* after hydrogen peroxide treatment and in *C. glabrata* after Amphotericin B exposure. While CFU is more sensitive at lower stress levels, the SYTO 9/PI staining effectively distinguishes sublethal from lethal doses, offering a valuable alternative for rapid, high-throughput survival quantification.

## Introduction

The budding yeast subphylum (Saccharomycotina) contains many important biological and biotechnological models as well as important human fungal pathogens, including *Candida albicans* [1], *Candida glabrata* [2], and the newly emerged *Candida auris* [3]. A common and critical task in the studies of this group, including both biology and biomedical research [4] as well as in environmental monitoring [5] and the food industry [6], is to quantify survival after stress or drug-treatment. The most widely-used and accepted method is the Colony Forming Unit (CFU) assay, which is relatively easy to perform and robust, but can be labor-intensive when assaying large numbers of samples and time-consuming due to the need to wait for single colonies to form. Therefore, there is a need for alternative methods that are fast and scalable for quantifying post-stress survival in this group.

There are two common measures used to quantify post-stress or post-drug treatment outcomes. The first one is represented by CFU, which measures viability, or the ability of cells to survive and reproduce to generate single colonies. The second is survival, or the lack thereof, i.e., cell death. Viability and survival (or cell death) measure related but distinct quantities: cells that are alive after the stress treatment could either die later or stay in a non-proliferative state, such as senescence [7], such that they do not contribute to the growth quantified by CFU. Conversely, CFU can be influenced by the additional stress introduced when performing dilutions and plating, and also the media and culturing conditions, which may lead to underestimates of the number of viable cells. Adding to the complication is the lack of a universal definition, hence methods to quantify cell death [8,9]. In this study, we follow the Nomenclature Committee on Cell Death and define cell death as the irreversible loss of plasma membrane integrity [9,10]. In yeast, a common way of monitoring plasma membrane integrity relies on the differential ability of certain dye molecules to penetrate intact *vs.* compromised plasma membranes, leading to a number of LIVE/DEAD assay reagents, most of which fluoresces upon binding nucleic acids [11].

Currently available assays to quantify post-stress outcomes present many limitations. CFU is robust and easy to perform, and its results are generally comparable between labs and studies. However, it is time consuming and labor intensive when assessing multiple samples. In a hypothetical experiment involving four samples with three replicates, to achieve ∼30-500 individual colonies per plate, three 10x dilutions of each sample are needed, resulting in nine plates per sample [12]. For *S. cerevisiae*, typically 48 hours of incubation is required for individual colonies to grow to a sufficient size for reliable counting [13]. This time can be even longer for other species. While modified CFU protocols have enabled higher throughput and automated counting, they require special setup or optimization, which limited their wide adoption [14–16]. As a result, manual counting of CFU is still a routine task performed in many microbiology labs.

Fluorescence dye-based methods are fast – mostly requiring 10-30 minutes of incubation. Coupled with flow cytometry, they provide fast, scalable and single-cell level information on survival. However, their application is significantly limited by a lack of standardized protocols and no systematic comparison with CFU results. In the literature, live/dead staining are nearly always used with microscopy to provide a qualitative representation of the survival proportion rather than yielding quantitative information [17]. In the few studies where quantification was performed, method details were scarce and varied [18,19].

Our goal in this study is to systematically characterize and optimize a fluorescence dye-based LIVE/DEAD assay for quantifying post-stress outcome in yeasts. We chose a two-dye kit composed of SYTO 9 and propidium iodide (PI). SYTO 9 labels all cells, live or dead, while PI is excluded from live cells with intact plasma membranes, hence differentiating between them. Moreover, the excitation and emission spectra of the two dyes make them an efficient pair for Förster Resonance Energy Transfer (FRET, Fig. 1A). FRET plus competitive exclusion of SYTO 9 by PI, whose affinity for DNA is higher [20], lead to a strong red fluorescence and muted green emissions in dead cells compared with live cells, hence better separating the two populations than using PI alone (Fig. 1B). This combo was initially developed for quantifying bacterial survival in environmental samples, known as BacLight [21]. Later it was applied to the baker’s yeast *S. cerevisiae* [19] and marketed as “FungaLight”. Despite its potential as a fast and scalable alternative for survival quantification, its use in the yeast literature is spotty and limited to mostly qualitative assessment. For example, when FungaLight was used to quantify the effect of antifungals on *C. albicans* survival, results were presented as microscopy images and were not quantified [22–29]. In the few studies where samples were assayed with flow cytometry, critical information such as the staining procedure, flow cytometry settings and gating strategies were often missing, making it difficult to compare or reproduce their results [18,19,30]. We believe the lack of systematic characterization of the method and standardization of the protocol is a major factor limiting the use of STYO 9/PI in quantitative assessment of post-stress survival.

**Fig. 1.**
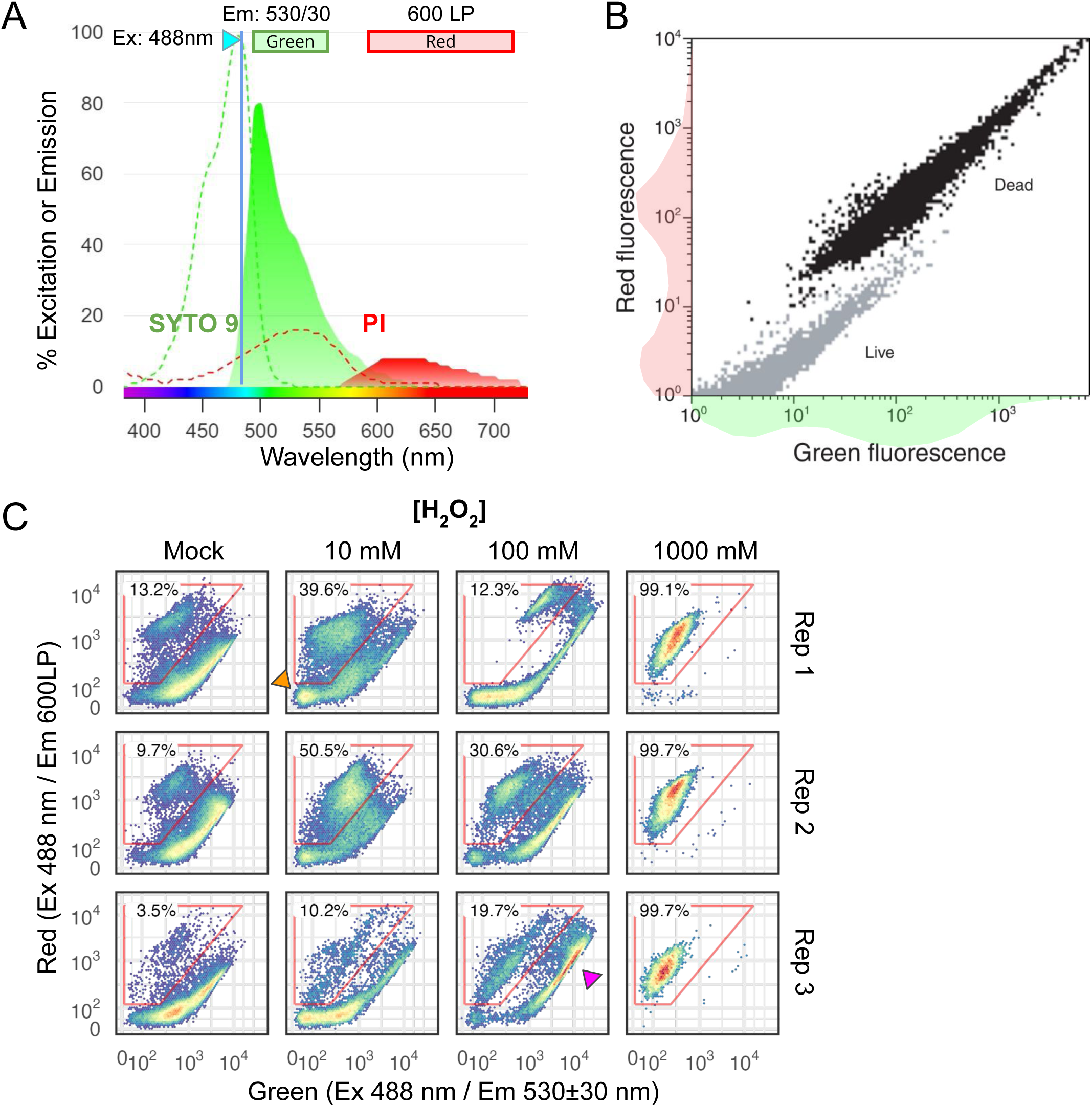
SYTO 9/PI (FungaLight) LIVE/DEAD stain, properties and application to H_2_O_2_-treated *C. glabrata* cells. (A) Excitation and emission spectra for SYTO 9 and Propidium Iodide (PI) when bound to DNA. Dashed lines and filled curves show the excitation and emission spectra, respectively. Both are normalized to the 488 nm laser. The plot is generated using FluoroFinder’s Spectra Viewer. (B) 2D density plot showing the dead (black) and live (gray) populations, reproduced from the FungaLight manual with permission. 1D density graphs shown next to the x- and y-axes. **(C)** 2D density plots for *C. glabrata* treated with different doses of H_2_O_2_ and stained with FungaLight. The red polygon gates identify dead and dead-like cells, with percentages on the top. A magenta arrowhead points to a subpopulation distinct from either live or dead cells as shown in B. Three biological replicates are shown as rows.

In this study, we systematically optimized the SYTO 9/PI LIVE/DEAD stain for use with flow cytometry to quantify post-stress survival in diverse yeast species. We found distinct staining patterns in cells that were damaged from those that were dead. We applied the optimized protocol to quantify the survival of several yeast species after stress or drug treatments, showing its broad applicability.

## RESULTS

### SYTO 9/PI staining for H_2_O_2_-treated *C. glabrata* cells show complex patterns and high sample-to-sample variability

We applied SYTO 9/PI staining on *C. glabrata* cells treated with different doses of H_2_O_2_ from 0 mM (mock) to a lethal dose of 1 M for 2 hours. We followed the manufacturer’s protocol for FungaLight and recorded the fluorescence signals in the green (530±30 nm) and red (600 nm long pass, or LP) channels with a blue laser (488 nm) as the excitation (Materials and Methods). At the highest H_2_O_2_ dose, all cells exhibited low green and high red fluorescence as expected for cells with a compromised plasma membrane (Fig. 1C). In mock-treated samples, the majority of the cells exhibited higher green than red signals, consistent with the expected patterns for live cells. However, we observed a subpopulation (red polygon gate) that exhibited “dead-like cell” features: they had lower green and high red signals, with the red fluorescence intensity comparable with or exceeding that of the dead cells (1 M H_2_O_2_-treated). The fraction of this subpopulation varied between 3.5%-13.2% across three biological replicates. This was unexpected given that the cells were taken from an exponentially growing population. Indeed, when we measured the CFU of the mock-treated sample and compared it to the number of cells plated (estimated by flow cytometry), we obtained a nearly 1:1 ratio (Fig. S1).

Compared with the mock, we observed an increase in the “dead-like cell” frequency in the 10 mM H_2_O_2_-treated samples, varying from 10% to 50% across replicates. There was also a subset of cells with very low red and green signals (orange arrowhead), which we hypothesize are unstained cells. At the severe 100 mM H_2_O_2_ dose, rather than a further increase in the fraction of “dead-like cell” population, we observed an increase in the population of cells with high-red and high-green signals (magenta arrowhead). Overall, the patterns we observed for H_2_O_2_-treated *C. glabrata* cells were more complex than what was shown in the FungaLight manual for a mixture of live and dead cells (Fig. 1B). There were also large sample-to-sample variations that prevented the assay from being reliably used to quantify post-stress survival outcomes. In the sections that follow, we systematically tested different parts of the staining protocol to improve the reproducibility and resolve some of the unexpected patterns.

### Using 0.85% saline as the staining buffer minimizes the dead-like and unstained cell populations

The FungaLight manual recommended washing and resuspending the cells in “appropriate buffer” and didn’t discuss the potential effects of the buffer on the staining pattern. In our initial test, we washed and resuspended cells in room temperature deionized water. To test if staining buffer choice affects staining results, we compared four buffers, i.e., deionized water, Phosphate Saline Buffer (PBS), 0.85% saline solution and Synthetic Defined (SD) media, the last being the base media in our experiments. The same mock- or H_2_O_2_-treated cultures were split four-ways, washed, resuspended and stained in the respective buffer, and the staining results were compared (Fig. 2A). We found that deionized water resulted in a large amount of “dead-like” staining events (∼60%, Fig. 2B top). SD media resulted in the lowest amount of “dead-like” cells but also produced far more unstained cells than the other three buffers (Fig. 2B, lower). By comparison, 1xPBS and 0.85% saline solution both provided a low amount of “dead-like” and unstained cells (Fig. 2B), supporting their use as the buffer for FungaLight staining. In our test, 0.85% saline performed slightly better than PBS. Hence, we use 0.85% saline for all subsequent experiments.

**Fig. 2.**
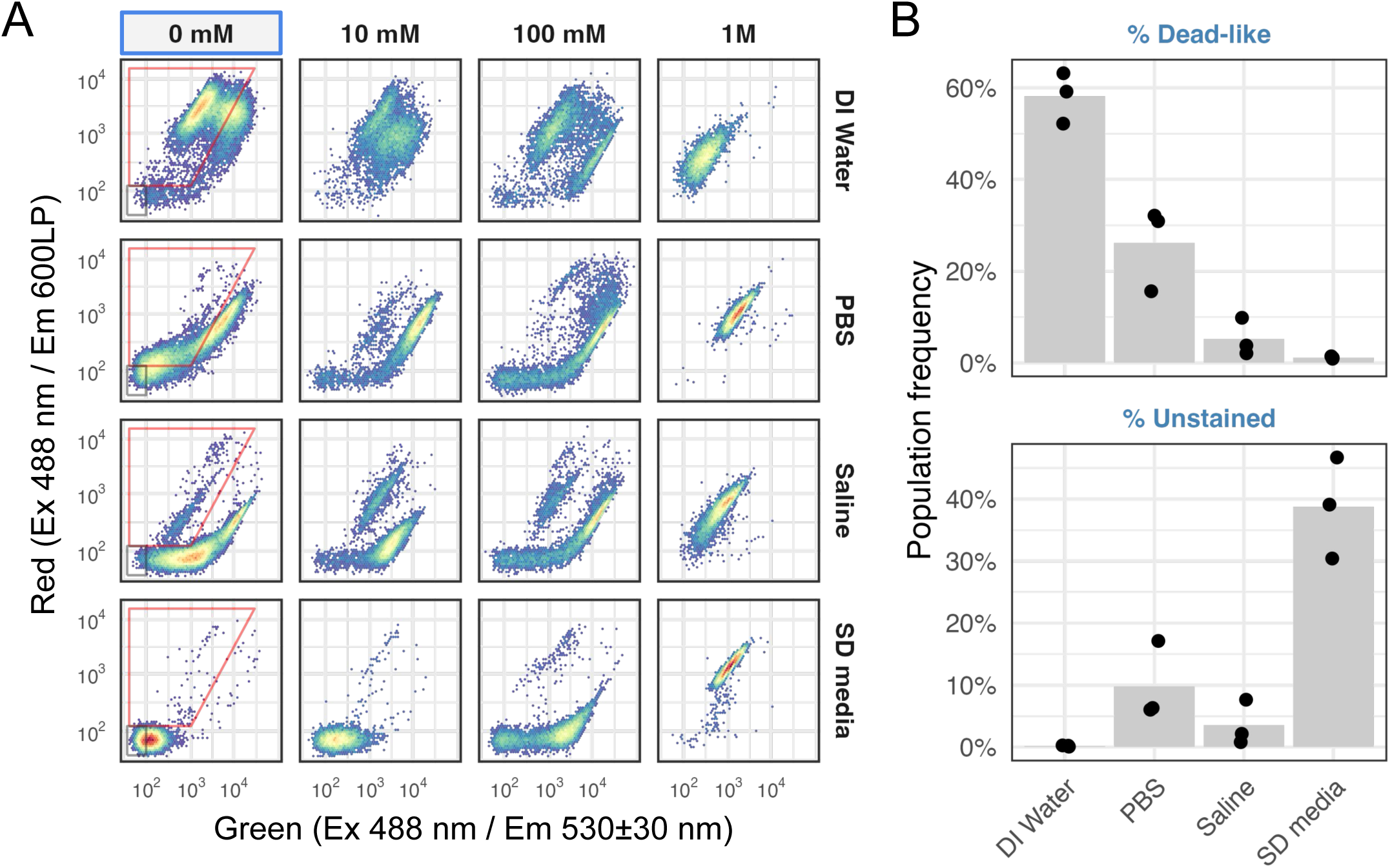
Effect of the staining buffer choice on the dead-like and unstained populations in mock-treated and FungaLight stained samples. (A) 2D density plots showing the FungaLight staining patterns of *C. glabrata* cells treated with different doses of H_2_O_2_, then stained in one of the four buffers: deionized water, 1x PBS, 0.85% Saline and Synthetically Defined (SD) media. In the mock-treated samples, the red trapezoid gate identifies the “Dead-like” population while the gray rectangular gate identifies the “Unstained” population. Their frequencies are further analyzed below. (B) Frequencies of the dead-like and unstained populations in the mock-treated samples (n = 3, biological replicates). Bars show the mean and dots show the individual replicates.

### Optimizing dye concentrations, staining time, and dye ratios

We then investigated the concentration and staining time for each dye individually as potential factors that could influence the result. The manual recommended using SYTO 9 and PI at 1000x dilutions from the 3.34 mM and 20 mM stock solutions, respectively. For both dyes, we tested 250x, 500x, 1000x and 2000x dilutions. For SYTO 9, we found that at 500x and 250x dilutions, more cells in the mock-treated sample fell within the “high-red and high-green” population seen in the 100 mM H_2_O_2_-treated sample (19% and 70%, respectively, Fig. S2A). Conversely, a 2000x dilution resulted in >90% events with a green fluorescence less than 10^2^, suggesting they were not stained. We therefore conclude that 3.34 uM (1000x dilution) is optimal for SYTO 9. For PI, we found minimal variation at the four dilutions tested for the mock-treated sample (Fig. S2A).

Fluorescence dyes take time to penetrate cells and bind to their target molecules, after which their signal could decay over time due to instability and photobleaching [31]. The manual recommended incubating the sample with the dyes for 15-30 minutes at 37°C and protected from light. We investigated changes in the SYTO 9 and PI’s signal intensity as a function of time when incubated at room temperature in the dark. Using exponential phase *C. glabrata* cells, we found that SYTO 9 signals plateaued at 15 min and didn’t change significantly up until 55 min post-staining, suggesting that an incubation time between 15-50 minutes can be used for SYTO 9 (Fig. S2B). Because PI is not permeable to live cells, we used heat-killed *C. glabrata* cells and found that the PI signal intensity reached a plateau within 5 minutes and began to decrease after 35 minutes (Fig. S2C). Together, we found the recommended 15-30 minutes of staining in the dark to be optimal for the combination of the two dyes.

We also tested different PI-to-SYTO 9 ratios, reasoning that the relative concentration of PI to SYTO 9 could affect the competitive exclusion and FRET between the two. However, we didn’t find significant differences in the staining pattern between 2:1, 1:1 and 1:2 ratios (1000x dilutions used as the base level) (Fig. S2D).

In summary, our systematic tests suggest that PI and SYTO 9 should be used at a 1000x dilution as the manual recommends; samples should be incubated with both dyes in the dark for 15 to 30 minutes at room temperature for both dyes to reach their stable signal intensity and assayed within 45 minutes.

### Sublethal dose of oxidative stress results in enhanced SYTO 9 staining but not PI

In our initial test of SYTO 9/Pi staining on H_2_O_2_-treated *C. glabrata* cells, we observed an unexpected population of cells with a high-green and high-red fluorescence (Fig. 1C, magenta arrowhead). This population persisted after we optimized the protocol, suggesting it is not a staining artifact. We noted that cells in this population showed stronger fluorescence than live cells in both channels, and they are distinct from dead cells, which have enhanced red and mute green fluorescence due to FRET and competitive exclusion of SYTO 9 by PI. Because SYTO 9 fluorescence is stronger than PI at the experimental conditions [20], and its emission bleeds into the red channel (Fig. 1A), we hypothesize the high-green and high-red fluorescence population contain cells with a higher level of SYTO 9 than live cells but little PI. Furthermore, because this population is more prominent in the 100 mM H_2_O_2_-treated sample, we hypothesize that severe, sublethal doses of H_2_O_2_ resulted in a population of oxidatively damaged cells that are more permeable to SYTO 9 but not yet permeable to PI. This hypothesis is consistent with the finding that in some bacterial species dead cells accumulate more SYTO 9 than live cells [31].

To test the hypothesis that the high-green and high-red signals are due to an increased SYTO 9 level, we performed single-dye staining for *C. glabrata* cells treated with increasing doses of H_2_O_2_ and quantified their emission patterns in both the green and red channels (Fig. 3A). Consistent with our hypothesis, SYTO 9-stained cells showed increased fluorescence when the concentration of hydrogen peroxide increased (Fig. 3A). In the mock sample, only ∼1% of the cells had a red fluorescence level greater than 10^2^; this proportion jumped to ∼25% and ∼50% in the 100 mM and 1 M H_2_O_2_ treated samples (Fig. 3B, Holm-Bonferroni corrected Student’s t-test *P* < 0.05 when compared with the mock). By comparison, cells stained with PI alone only showed a significant increase in the proportion of “PI-positive” cells at the lethal 1 M H_2_O_2_ dose (Fig. 3B, Holm-Bonferroni corrected Student’s t-test *P* < 0.5 compared to the mock). We also calculated the median fluorescence intensity (MFI) for all cell events in the SYTO 9 stained samples, confirming that the MFI increased significantly in both green and red channels at the 100 mM and 1 M H_2_O_2_ doses (Fig. 3C). We conclude that at the sublethal dose of 100 mM H_2_O_2_, ∼25% of the *C. glabrata* cells became more permeable to SYTO 9 but not yet to PI. This likely leads to the “Intermediate” high-green and high-red subpopulation in the SYTO 9/PI staining (Fig. 1C). At the lethal 1 M dose, however, cells become more permeable to both SYTO 9 and PI. Competitive exclusion of SYTO 9 by PI and FRET between the two result in enhanced red and reduced green fluorescence relative to live and the “Intermediate” population.

**Fig. 3.**
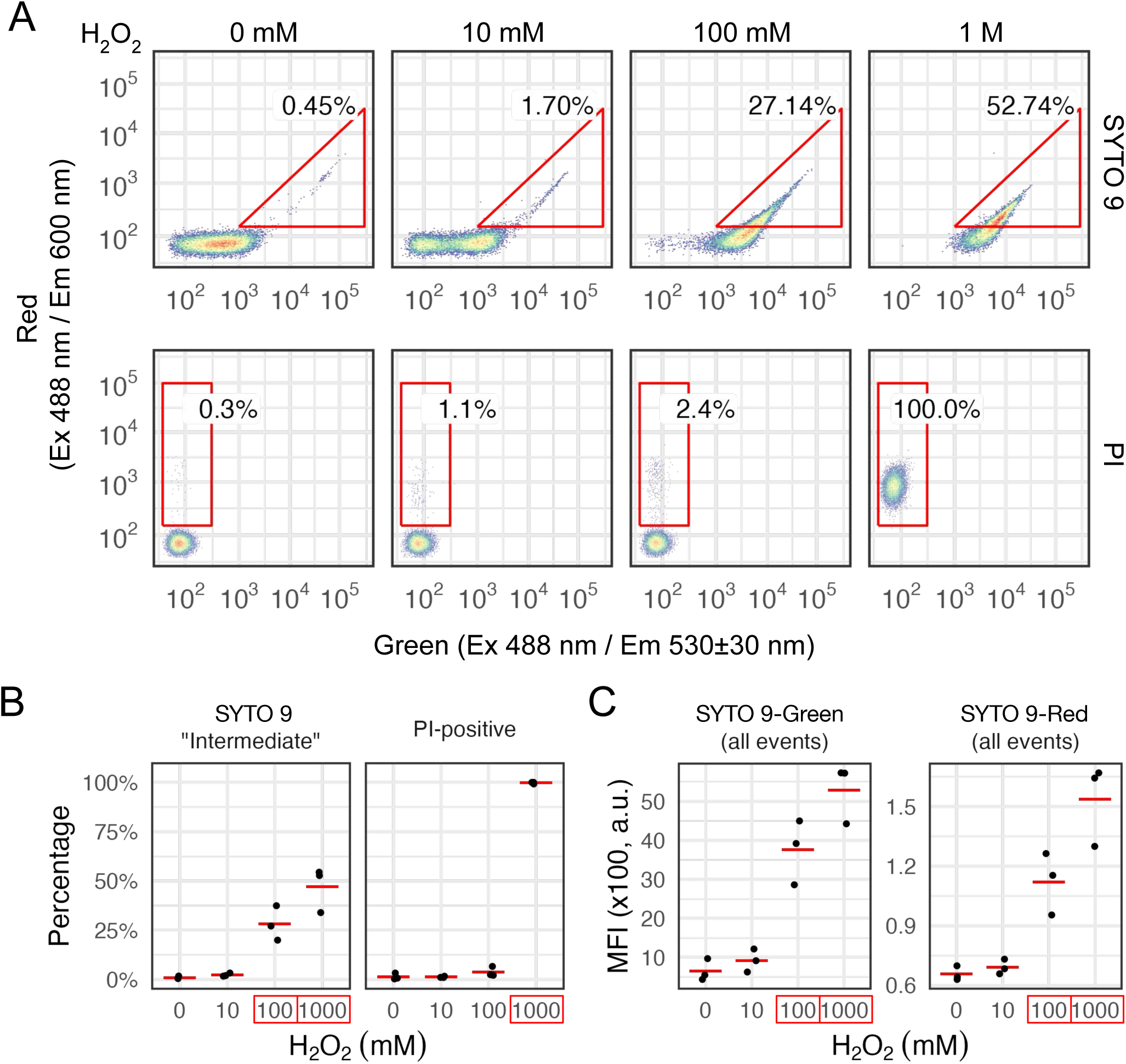
*C. glabrata* cells exposed to sublethal doses of H_2_O_2_ accumulate more SYTO 9 but not PI. (A) 2D density plots showing *C. glabrata* cells treated with various doses of H_2_O_2_ for 2 hrs and stained with either SYTO 9 or PI individually. An “intermediate” high-green and high-red population is shown with a triangle gate for the SYTO 9 stained samples (top) and a “PI-positive” population shown with a rectangular gate for the PI stained samples (bottom). Percentages of gated events are shown next to the gates. (B) Percentages of the SYTO 9 “Intermediate” and PI-positive populations are shown as dots and their means as red lines (n = 3, biological replicates). A significant difference compared to the mock-treated (0 mM) sample was determined by a linear model. A Holm-Bonferroni corrected Student’s t-test P-value < 0.05 is indicated by a red box around the x-axis label. (C) Median Fluorescence Intensity (MFI) for all events in samples stained with SYTO 9 are shown for both the green and the red channels. Biological replicates are shown as dots and their means as red lines. Significant differences from the mock-treated sample were determined and indicated the same way as in (B).

### SYTO 9/PI is less sensitive than CFU at medium levels of H_2_O_2_ but can better distinguish sublethal from lethal H_2_O_2_ doses

Having optimized the staining protocol for SYTO 9/PI (referred to as FungaLight from hereon), we next compared its result to two widely cited methods for quantifying post-stress or drug treatment outcomes, i.e., CFU and PI staining. The former is generally accepted as the standard method for viability while the latter is frequently used to quantify live/death based on plasma membrane permeability [32].

To compare these methods, we treated *C. glabrata* cells with 0, 10, 100, and 1000 mM of H_2_O_2_ for 2 hours. The same post-treatment sample was used for CFU as well as PI or FungaLight staining in separate vials. The percent viable/live cells were quantified and compared. For PI, we gated for live cells only; for FungaLight, we additionally gated for the “Intermediate” population, which we hypothesize contains oxidatively damaged cells that are more permeable to SYTO 9 but not yet permeable to PI (Fig. 4A).

**Fig. 4.**
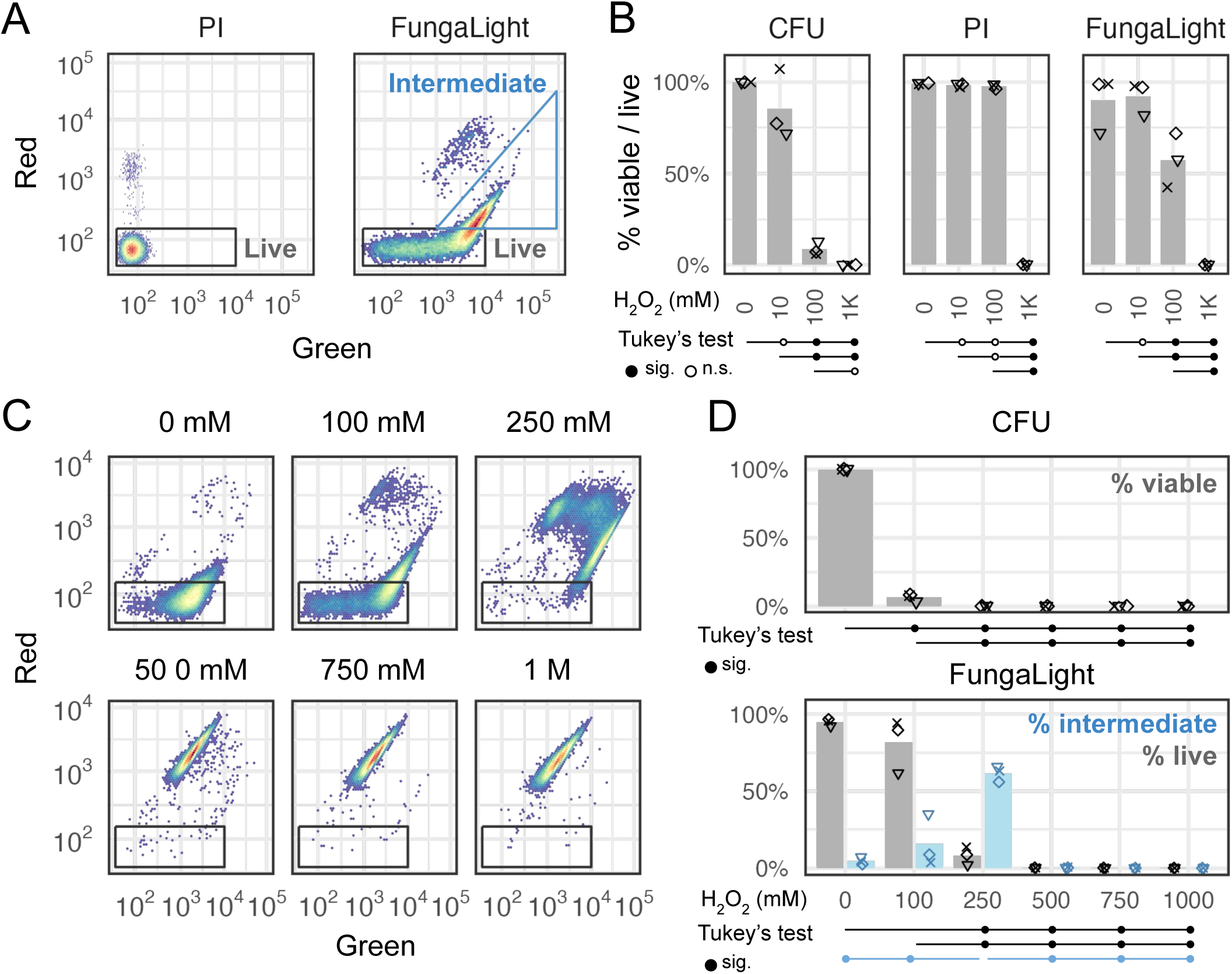
Comparing CFU, Propidium Iodide (PI), and FungaLight in the estimates of viability or survival. (A) Gating strategy for PI and FungaLight stained samples (see Methods for gate boundary coordinates). (B) C. glabrata cells were treated with H_2_O_2_ at various concentrations for 2 hours and then plated for CFU or stained with PI alone or with FungaLight. Percent viable (for CFU) or percent live (PI and FungaLight) estimates for three replicates were shown as dots (n = 3, biological replicates). Bars are the means of the replicates. Dot shapes distinguish different days of experiments. Tukey’s range test (aka Tukey’s HSD) was used to compare all pairs of samples. The results are presented below the x-axis: lines originate from the first sample and end with the other sample, with a filled circle at the end indicating a significant difference at 0.05 level, or open circle otherwise (n.s. = not significant). (C) Density plots for one replicate showing the FungaLight staining patterns for *C. glabrata* cells treated with H_2_O_2_ from 0 to 1 M in 100 mM increments. The axes and live cell gates were the same as in A. Intermediate gates were omitted for clarity. (D) CFU and FungaLight survival estimates for the same concentration series of H_2_O_2_ in *C. glabrata* (n = 3, performed on three different days indicated by the shape of the dots). Bars represent the mean. For FungaLight, the gray bars and dots indicate the percent of events in the live gate and the light blue bars and dots show the percent of events in the “Intermediate” gate (see A). Tukey’s range test was applied to (i) CFU, (ii) FungaLight % live events and (iii) % intermediate events separately. Only significant results at a 0.05 level were indicated by lines and filled circles due to space constraints.

PI-staining detected a significant drop in the percent-live cells only at the lethal 1 M H_2_O_2_ dose (Fig. 4B, middle, Tukey’s range test *P* < 0.05). By contrast, CFU results revealed a significant decrease in viability at the 100 mM and 1M H_2_O_2_ doses compared with the mock and 10 mM H_2_O_2_-treated samples (Fig. 4B). While a drop in CFU in the 10 mM H_2_O_2_ treated sample was observed compared with the mock, the difference was not significant by Tukey’s range test (aka Tukey’s HSD) at the 0.05 level. The difference in CFU between the 100 mM and 1 M H_2_O_2_-treated samples was also insignificant (Fig. 4B).

Similar to CFU, FungaLight percent-live estimates also revealed a significant decrease in survival in the 100 mM and 1 M H_2_O_2_-treated samples compared with the mock and 10 mM H_2_O_2_-treated ones. However, FungaLight estimates for the 100 mM H_2_O_2_-treated sample were ∼50%, much higher than its CFU and the difference between this condition and the 1 M H_2_O_2_ treatment was significant (Fig. 4B, right). It is worth noting that the two methods measure different quantities and therefore their values are not directly comparable – CFU quantifies the ability of a cell to recover from the oxidative stress and reproduce to form a colony 48-72 hours post treatment, while FungaLight measures the plasma membrane permeability immediately after the treatment. Nonetheless, the goal of our study is to compare these two methods at a practical level for quantifying post-stress outcomes.

Since FungaLight revealed a significant difference between the 100 mM and 1M H_2_O_2_-treated samples, we asked if it is able to resolve different levels of damage caused by a gradient of sublethal doses of oxidative stress. To test this, we treated *C. glabrata* cells with a linear gradient of H_2_O_2_ from 100 mM to 1 M, then subjected the treated samples to both CFU and FungaLight staining. For the latter, 2D density plots showed distinct fluorescence patterns between the mock, 100 mM and 250 mM H_2_O_2_-treated samples; above 250 mM, all events appeared in the “dead” population (Fig. 4C). Quantitatively, we found that CFU was able to detect a significant difference between all non-zero doses of H_2_O_2_ vs the mock sample as well as a significant difference between the 100 mM dose against all higher doses by Tukey’s range test at a 0.05 level (Fig. 4D top, only significant tests were labeled below the plot). FungaLight detected a significant difference between doses at 250 mM or above vs either the mock or 100 mM H_2_O_2_-treated sample, but not between the 100 mM H_2_O_2_-treate and the mock sample (Fig. 4D bottom). In general, we observed a higher sample-to-sample variation in the FungaLight results than CFU, limiting its power to distinguish post-stress survival especially at a lower dose. However, we noticed that the percent of “Intermediate” population was significantly different between the 250 mM H_2_O_2_ dose and all other doses. This extra dimension offers additional information and provides more gradation in assessing the outcomes of oxidative stress.

In summary, we found that PI-staining only detects changes when a lethal H_2_O_2_ dose of 1 M was used. Notably, PI estimated percent-live values for 100 mM H_2_O_2_-treated samples were close to 100% even though CFU shows <10% of the cells were viable following the treatment. Compared with CFU, FungaLight (SYTO 9/PI) can similarly differentiate the sublethal dose of 100 mM from the mock and mild 10 mM dose. While it exhibited a higher sample-to-sample variance, limiting its power in detecting differences at a lower dose of H_2_O_2_, it revealed more information through changes in the percent “Intermediate” population at 100 mM and 250 mM H_2_O_2_ doses. These observations suggest that FungaLight can be a useful alternative to CFU for quantifying post stress survival especially at higher, sublethal doses.

### FungaLight can be used to quantify post-stress or drug treatment survival across diverse yeast species

To determine if the optimized FungaLight assay works in other yeast species, we applied it to H_2_O_2_-treated *S. cerevisiae* and *C. albicans* cells. We chose these two species as they are both the most important and frequently studied yeasts and also at different evolutionary distance from *C. glabrata*: the baker’s yeast *S. cerevisiae* diverged from *C. glabrata* ∼80 million years ago post the Whole Genome Duplication (WGD) event while *C. albicans* diverged from both *S. cerevisiae* and *C. glabrata* approximately 240 million years ago prior to the WGD (Shen et al 2018). We used different concentrations of H_2_O_2_ for the two species based on our preliminary experiments with the goal of selecting a serial dilution series of the oxidant to achieve comparable CFU-based viability as compared to our results for *C. glabrata*. Despite the literature reporting a higher H_2_O_2_ resistance in *C. albicans* compared to *S. cerevisiae* [33], we found similar resistance levels in the two species, both lower than that of *C. glabrata*. We chose 0.6 and 6 mM in place of 10 and 100 mM as did in *C. glabrata*. We kept the 1 M as the lethal dose.

We applied the same FungaLight protocol to both species after 2 hours of H_2_O_2_ treatment. The fluorescence patterns in the red and green channels are very similar to what we found in *C. glabrata*. However, the absolute intensities shifted between species. We manually adjusted the live and intermediate cell gates for each of the two species to capture the same population based on visual inspections (Fig. 5A, Methods). We also performed CFU assays on the same samples as we did FungaLight staining with, then compared CFU with FungaLight percent live and percent intermediate values (Fig. 5B, C). In both species, CFU estimates were significantly different between the mock and all three H_2_O_2_-treated samples as well as between different doses of H_2_O_2_ except for the 6 mM vs 1 M comparison (Fig. 5B, C, top). FungaLight percent live values were much higher in the 0.6 and 6 mM H_2_O_2_-treated groups than CFU in both species. In *S. cerevisiae*, significant differences were detected in all pairwise comparisons for the percent live values except for the mock vs 0.6 mM pair (Fig. 5B, bottom). In *C. albicans*, while the general trend was similar, a higher sample variance resulted in non-significant differences for most of the percent live value comparisons except for those involving the 1 M H_2_O_2_-treated group (Fig. 5C, bottom). Percent intermediate values were the highest in the 6 mM group, which were significantly higher than the rest in *S. cerevisiae*, but not in *C. albicans* (Fig. 5B, C, bottom). These results showed that the FungaLight assay can be used to quantify post-oxidative stress survival in diverse yeast species; its results are consistent in ranks with CFU. Similar to our observation in *C. glabrata*, FungaLight can have a higher sample-to-sample variance, but it is useful in distinguishing between medium to lethal doses while CFU is more sensitive towards lower doses. Also, we note that while the same protocol is applicable across multiple species, the percent live and intermediate gate boundaries need to be adjusted for each species based on their mock-treated and dead cell controls.

**Fig. 5.**
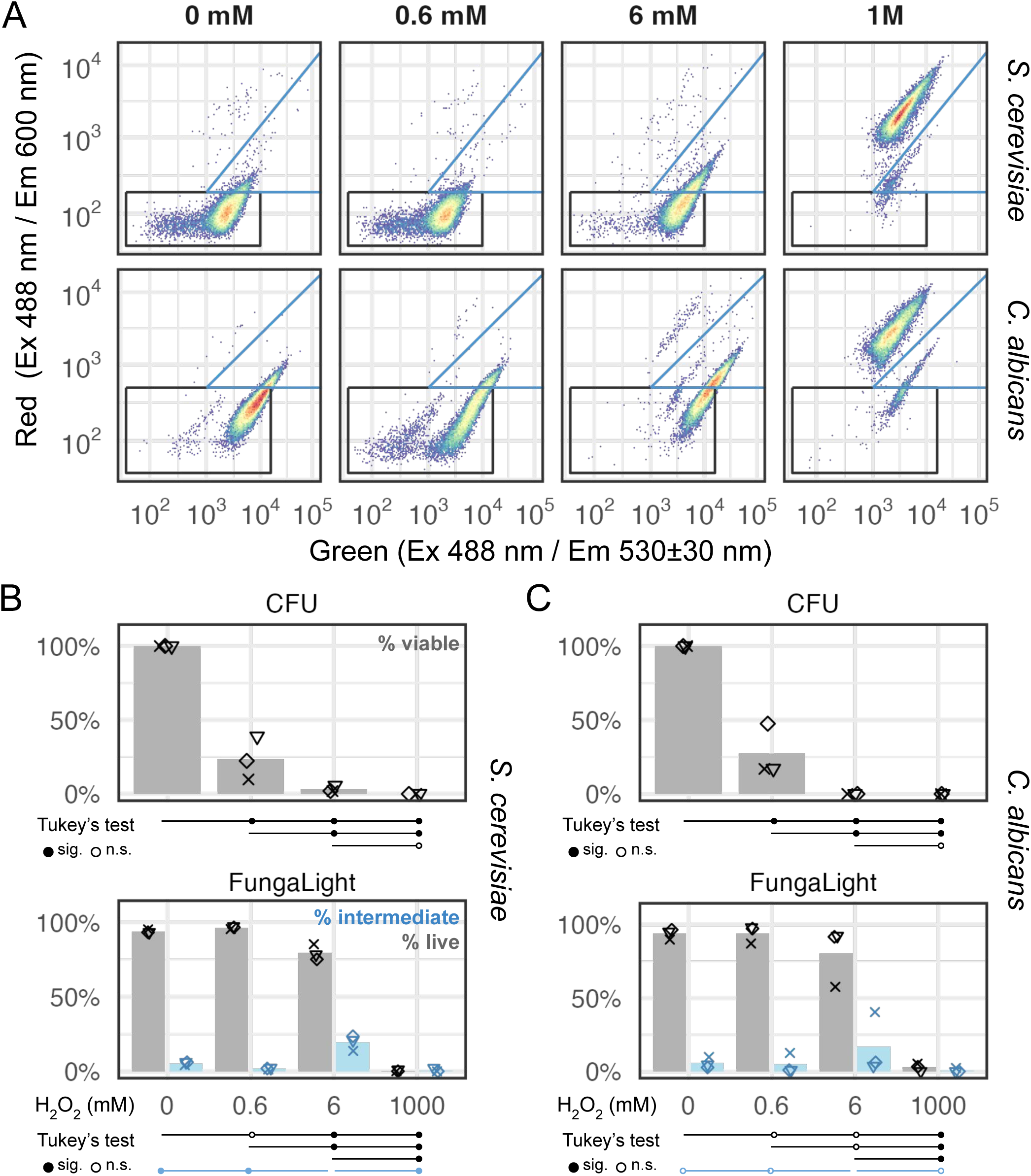
FungaLight quantifies post-oxidative stress survival in diverse yeast species. (A) Representative 2D density plot for *S. cerevisiae* and *C. albicans* cells treated with different doses of H_2_O_2_. The live (black) and intermediate (blue) cell gates were shown (see Methods for gate boundary coordinates). (B) Percent viable (for CFU) or percent live (FungaLight) estimates for *S. cerevisiae* cells treated with 0 to 1 M H_2_O_2_ (2 hours) were shown as dots (n = 3, biological replicates). Bars represent the means. Dots with the same shape represent the same day of experiment across both methods. Tukey’s range test (aka Tukey’s HSD) was used to compare all pairs of samples. Test results were presented below the x-axis in the same way as in Fig. 4. (C) Same as B for *C. albicans* samples.

To determine if FungaLight can be used to quantify post-treatment outcomes beyond the oxidative stress, we applied the same protocol to *C. glabrata* cells treated with Amphotericin B (AmpB), a frontline antifungal used to treat systemic candidiasis [34]. We observed a similar “live → intermediate → dead” shift in the FungaLight staining pattern with increasing doses of AmpB from 0 to 1 ng/mL, showing the applicability of FungaLight in this application (Fig. 6A). Viability estimates based on CFU showed a roughly linear decrease with an increasing AmpB dose (Fig. 6B). Significant differences were determined by Tukey’s range test and shown at the bottom of Fig. 6B. FungaLight estimates showed a larger sample variance. Percent live events showed significant differences only between the two or three highest doses when compared with the mock or the 0.04 ng/mL dose (Fig. 6C). Percent intermediate values were markedly higher in the 0.08 and 0.1 ng/mL AmpB-treated groups: there were significant differences between these two groups and the 0.04 ng/mL or the 1 ng/mL-treated groups (Fig. 6C). Finally, we observed a strong linear correlation between the percent live values and CFU in the AmpB-treated *C. glabrata* cells (Fig. 6D, Pearson’s r = 0.94, Spearman’s ρ = 0.98).

**Fig. 6.**
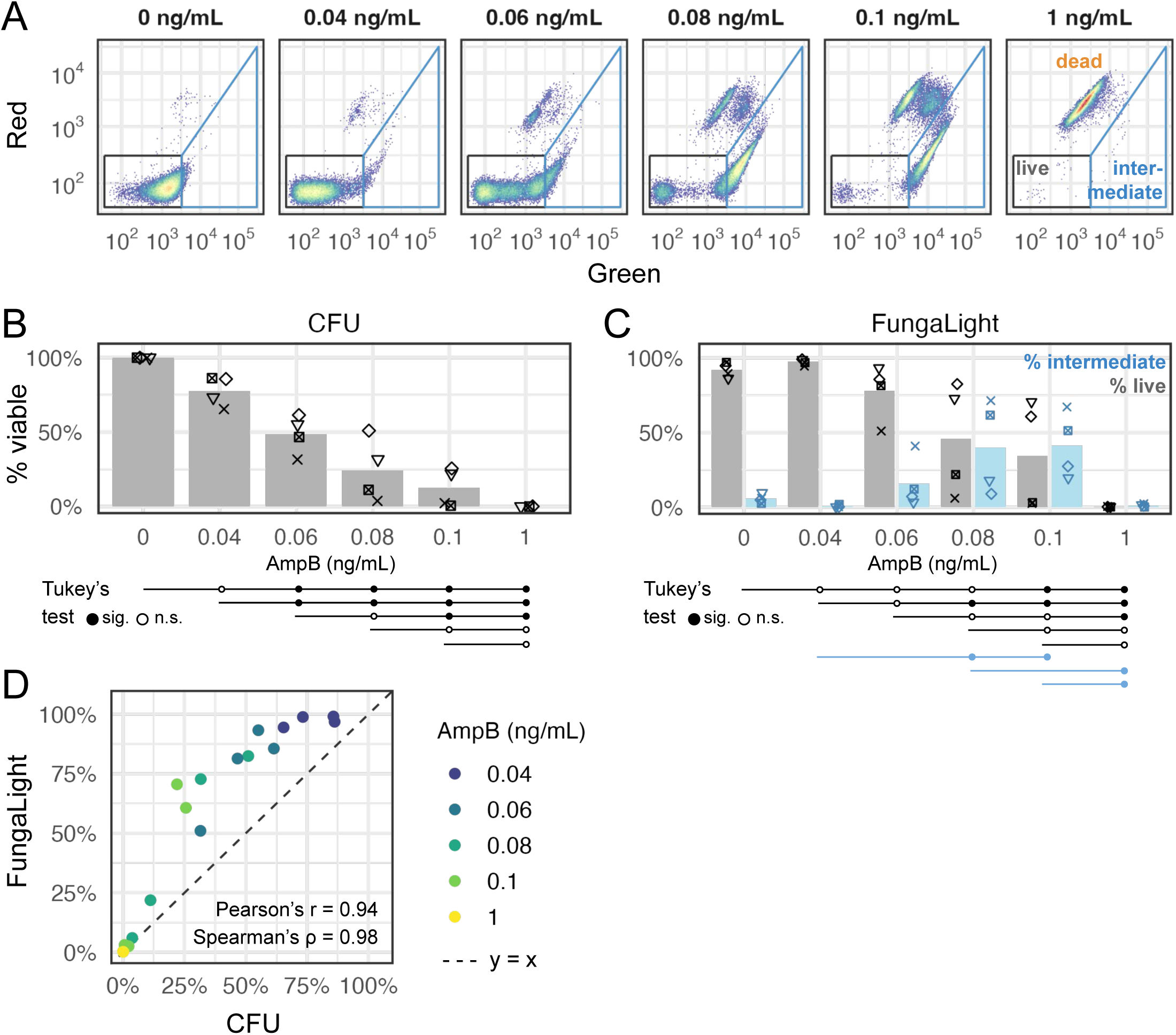
FungaLight quantifies survival after treatment with the antifungal Amphotericin B (AmpB). (A) Representative 2D density plot for *C. glabrata* cells treated with different doses of AmpB. The live (black) and intermediate (blue) cell gates were shown (see Methods for gate boundary coordinates). (B) Percent viable estimates for *C. glabrata* cells treated with 0 - 1 ng/mL AmpB shown as dots (n = 4, biological replicates). Bars represent the means. Dots with the same shape represent the same day of experiment. Tukey’s range test (aka Tukey’s HSD) was used to compare all pairs of samples. Test results were presented below the x-axis in the same way as in Fig. 4. (C) Same as B for FungaLight percent live and percent intermediate gated events. Only significant results at a 0.05 level were shown for the % intermediate events. (D) Scatter plot shows the relationship between the FungaLight % live and CFU % viable estimates. Dots are colored by AmpB dose. The dash line indicates y = x. Pearson’s correlation coefficient and Spearman’s rank correlation coefficient are shown inside the plot.

In summary, we successfully applied the FungaLight assay to quantify post-oxidative stress survival in three yeast species, as well as post-antifungal survival in one of them, *C. glabrata*, demonstrating the broad applicability of the assay and our optimized protocol.

## Discussion

Quantifying post-stress and drug treatment outcomes is a routine yet critical task in studies of diverse yeast species for basic, medical and industrial purposes. In this study we systematically characterized the two-dye LIVE/DEAD assay: SYTO 9/PI, aka FungaLight. Through rigorous testing of buffer conditions, we found that 0.85% saline buffer minimizes staining artifacts such as unstained cells and “dead-like cell” populations, which were present in DI water and growth media-based staining buffer applications. We also determined the optimal dye concentrations and staining incubation time. Interestingly, we discovered that at sublethal doses of oxidative stress, an “Intermediate” cell population exhibited distinct fluorescence patterns than both the live and dead cell populations. Evidence supported our hypothesis that this population includes damaged cells with increased SYTO 9 uptake but minimal PI accumulation. This additional information enhanced the resolving power of FungaLight especially between medium to high doses of stress. Lastly, we applied the optimized FungaLight assay to two other yeasts and after an antifungal treatment, demonstrating its broad utility as a complementary method to traditional CFU assays.

Dye-based methods like FungaLight when coupled with flow cytometry are fast and scalable. Compared with CFU, which requires multiple dilutions and a lengthy incubation period for single colonies to emerge, a typical FungaLight assay can be done within 2 hours post stress or drug treatment. Using the autosampler attachment, we can conveniently examine a large number of samples in a 96-well format, further enhancing the scalability of the assay. Another advantage of FungaLight is that it reveals finer gradations in the damage to the cells caused by the stress or drug. Compared with CFU and single dye staining like propidium iodide (PI), which classifies cells into either live or dead, FungaLight staining reveals a transitional population of cells under sublethal stress levels, enhancing our ability to distinguish the outcomes at sublethal doses from those at lethal levels.

We also recognized several limitations of FungaLight in our characterization. The most important limitation is the need to manually assign the gates when analyzing the flow cytometry data for each new species and treatment condition. This is because the fluorescence signal intensity is influenced not only by the cell’s plasma membrane integrity, but also by staining and flow cytometry conditions as well as species-specific cell size and membrane permeability traits. The first group of variables, such as dye concentrations, flow cytometer voltage settings, can introduce sample-to-sample variations as seen in our experiments (Fig. 4-6), which reduces the statistical power for detecting differences between treatments and can make it difficult to compare results between different labs. The second group of species-specific variables mean that gate boundaries need to be calibrated for each new species and treatment, potentially restricting its wide adoption (Fig. 5, 6).

One possible source of sample-to-sample variance in FungaLight staining is variations in cell size distributions as a result of the stress treatment. It has been shown that H_2_O_2_ can cause cell size changes in a dose-dependent manner in cyanobacteria [35]. This, in turn, could affect the fluorescence intensity by the two dyes in FungaLight. We compared the distributions of the forward scatter height (FSC.H), which is correlated with cell size, among samples treated with different H_2_O_2_ doses. In all three species, we observed an interesting pattern where the low dose treatment group (10 mM for *C. glabrata* and 0.6 mM for *S. cerevisiae* and *C. albicans*) exhibited a right shift, indicating cell size expansion, while at medium to high doses, including the lethal 1 M dose, there was a slight leftward shift, indicating cell size contraction (Fig. S3). Normalizing the fluorescence signal by cell size (Materials and Methods), however, had only small effects on the variance and didn’t affect the statistical test results (Fig. S4).

Despite its limitations, our results showed that FungaLight is a valuable tool for yeast research and complements the traditional CFU. It is worth emphasizing, however, that the two assays measure fundamentally different traits: CFU quantifies viability while FungaLight like other dye-based methods measure plasma membrane integrity. Therefore, there can be genuine differences between the two measures for the same sample, such as when a subpopulation of cells enter senescence, which will not be classified as dead by FungaLight but also don’t contribute to CFU. Such differences are not a bug but a feature that should be appreciated and exploited. That being said, we imagine that in most practical applications, researchers want a method to quantify the effects of a stress or drug but do not explicitly distinguish viability from survival. In such cases, FungaLight can serve as a useful alternative to CFU to either achieve the results faster and / or to examine a large number of samples in parallel. We found the two methods generate broadly consistent results while differing in their resolving power at different parts of the stress level gradient. As shown for H_2_O_2_ and Amphotericin B-treated *C. glabrata* cells, CFU is better at differentiating mock and low dose treatments but lacks the ability to distinguish between sub-lethal vs lethal doses; FungaLight, by contrast, is more powerful in resolving differences between the sublethal and lethal doses, yet it doesn’t distinguish mock and low dose treatments (Figs. 4-6). Knowing these properties will help the researcher choose the proper method or combine their power.

In the future, we would like to improve the FungaLight assay by devising novel analysis methods that capture the essence of the observed fluorescence pattern, namely a gradual shift from the low-green, low-red “live” state to the medium-green, high-red “dead” state, through the “intermediate” high-green, high-red “damaged” state. A scale-free approach that doesn’t require manual gate drawing and thus not dependent on the absolute fluorescence scales will simplify the assay analysis while also making the results more comparable across studies. In summary, we believe our optimized FungaLight assay provides a rapid and scalable approach for quantifying yeast survival after stress and drug treatments. It can be used as an alternative to CFU when the researchers do not need to distinguish between viability and survival, or it can be used together with CFU to provide in-depth information for the post-stress states of the population.

## Materials and Methods

### Strains

Yeast species and strains used in this study are: *C. glabrata* BG99, derived from a clinical isolate, BG2 [36]; *S. cerevisiae* K699 (ATCC #200903); *C. albicans* SN152, derived from a clinical isolate, SC5314 [37]. All strains used are listed in Table 1.

**Table 1.** Yeast strains used in this study.

| Species name | ID | Genotype | Source |
| --- | --- | --- | --- |
| *C. glabrata* | yH181 | [BG2] his3Δ(1 + 631) | [36] |
| *S. cerevisiae* | yH154 | [W303] MATa ade2-1 trp1-1 can 1–100 leu2-3,112 his3-11,15ura3 GAL+ | ATCC #200903 |
| *C. albicans* | yH714 | [SC5314] SN152, HIS3 and LEU2 | [37] |

### Media and yeast culture

Yeast cultures were grown in Synthetic Complete (SC) media, made with Yeast Nitrogen Base without amino acids (Sigma Y0626), 2% glucose, and amino acid mix. For SC agar plates, 2.5% agar was added to the SC liquid media before autoclaving. SC agar plates were stored at 5 °C and used within two weeks. For hydrogen peroxide stress, a 30% (w/w, Sigma H1009) stock solution was added to the SC media to the indicated concentration and used within 30 minutes.

### Hydrogen peroxide treatment

For all three species of yeast, overnight culture was diluted in the morning in SC and grown to the mid-log phase (OD_600_ = 1). For each treatment, 600 μL of cells were transferred to a well in a 96 deep-well plate (Thermo Fisher 278606). The plate was centrifuged at 3,000g for 5 min and the media was removed by aspiration. Mock (SC) or hydrogen peroxide stress media was added to each well to reach an OD_600_ of 1. The plate was incubated for 120 min at 30 °C with shaking at 300 rpm.

### Colony Forming Units (CFU) assay

Yeast cells were collected by centrifugation at 3,000g for 5 minutes. After removing the media, cells were resuspended in 0.85% saline buffer and adjusted to an OD_600_ of 0.2. Serial dilutions were made to aim at 50-500 single colonies per plate (typically 100-1000 fold for H_2_O_2_-treated cells). 50 μL of the properly diluted sample was spread onto SC plates with sterilized glass beads. Plates were incubated at 30 °C for 48 hours. Colonies were tallied manually. Three biological replicates were used per treatment group.

### SYTO 9/PI, aka FungaLight staining

Dye preparation: LIVE/DEAD FungaLight Yeast Viability Kit for flow cytometry was obtained from Thermo Fisher (Cat# L34952). The kit contains two components, which we used either individually or in combination as indicated in the text. SYTO 9 (component A) was diluted 1:100 to a working concentration of 3.34 × 10^−2^ mM with deionized water immediately before staining. PI (component B) was diluted 1:100 to a working concentration of 20 × 10^−2^ mM with deionized water and stored for less than 6 months at 5 °C.

After mock or stress treatment, yeast cells were collected by centrifugation at 3,000 g for 5 minutes. The treatment media was removed by aspiration. Cells were resuspended in 0.85% saline buffer to an OD_600_ of 1 (except when testing different staining buffers, in which case another solution was used for resuspension).

To stain the sample, the working stock of SYTO 9 (3.34 × 10^−2^ mM) and PI (20 × 10^−2^ mM) were added at a 1:10 ratio unless stated otherwise. The final concentrations were 3.34 μM for SYTO 9 and 20 μM for PI. At least three biological replicates were performed per group.

To determine the time evolution of fluorescence signals for each dye, after either dye is added to the sample, fluorescence was measured every 5 min from 0 min at the time of adding the dye to 55 min after in the single dye time course. All incubations were performed in the dark at room temperature (∼25 °C) until being measured on the flow cytometer.

### Flow Cytometry

Samples were stained with one or both dyes of FungaLight. To quantify with flow cytometry, 20 μL of the stained sample was diluted into 200 μl of 0.85% saline buffer and directly measured on an Attune NxT instrument fitted with an autosampler (Thermo Fisher Cat#A24858). A voltage of 345 mV and 399 mV were used for the FSC and SSC channels.

For SYTO 9 staining alone, samples were excited at 488 nm and emission was collected using a 530+/-30 nm bandpass filter at 230 mV. For PI-staining alone, samples were excited by a 488 nm blue laser and emission was collected using a 600 nm longpass filter (>600 nm) at 200 mV. For PI and SYTO 9 co-staining, samples were excited at 488 nm, and emissions were collected in the BL1 channel using a 530+/-30 nm filter at 230 mV and in the BL3 channel using a 600 nm longpass filter (>600 nm) at 200 mV. Samples were run at 200 μl / min speed and at least 30,000 total events were collected per sample. At least three biological replicates were measured per treatment group.

### Flow cytometry data analysis

FCS (v3.1) files were exported from the Attune NxT software and analyzed in R using flowCore (2.12.0), openCyto (2.16.1), flowWorkspace (4.12.2), ggcyto (1.32.0) and tidyverse (2.0.0) packages in R. We used a standard gating strategy to gate for single cells: 1) Non-cell events were removed by gating on FSC.H and SSC.H (Fig. S5A); 2) single cell events were identified based on FSC.H and FSC.W (Fig. S5B). In the main text results, gated single-cell events were then plotted on the FungaLight Green (BL1-H) and FungaLight Red (BL3-H) axes. Live, intermediate and dead cell gates were specified for each species and treatment condition. The gate boundaries used are listed in Table 2. Percent live and intermediate gate events were calculated by dividing the number of events in each gate by the total number of single cell events. In Fig. S3 and Fig. S4 where we examined the effect of normalizing by cell size, the green channel fluorescence was divided by FSC.H^1.2 and the red channel fluorescence by FSC.H^0.8, then multiplied by a constant such that the resulting values have the same median as before the normalization. These normalized values became generally independent of FSC.H as a proxy for cell size. The same procedure as above was followed to calculate the percent live and intermediate events, except instead of the FungaLight Green and FungaLight Red fluorescence, the normalized values were used.

**Table 2.**
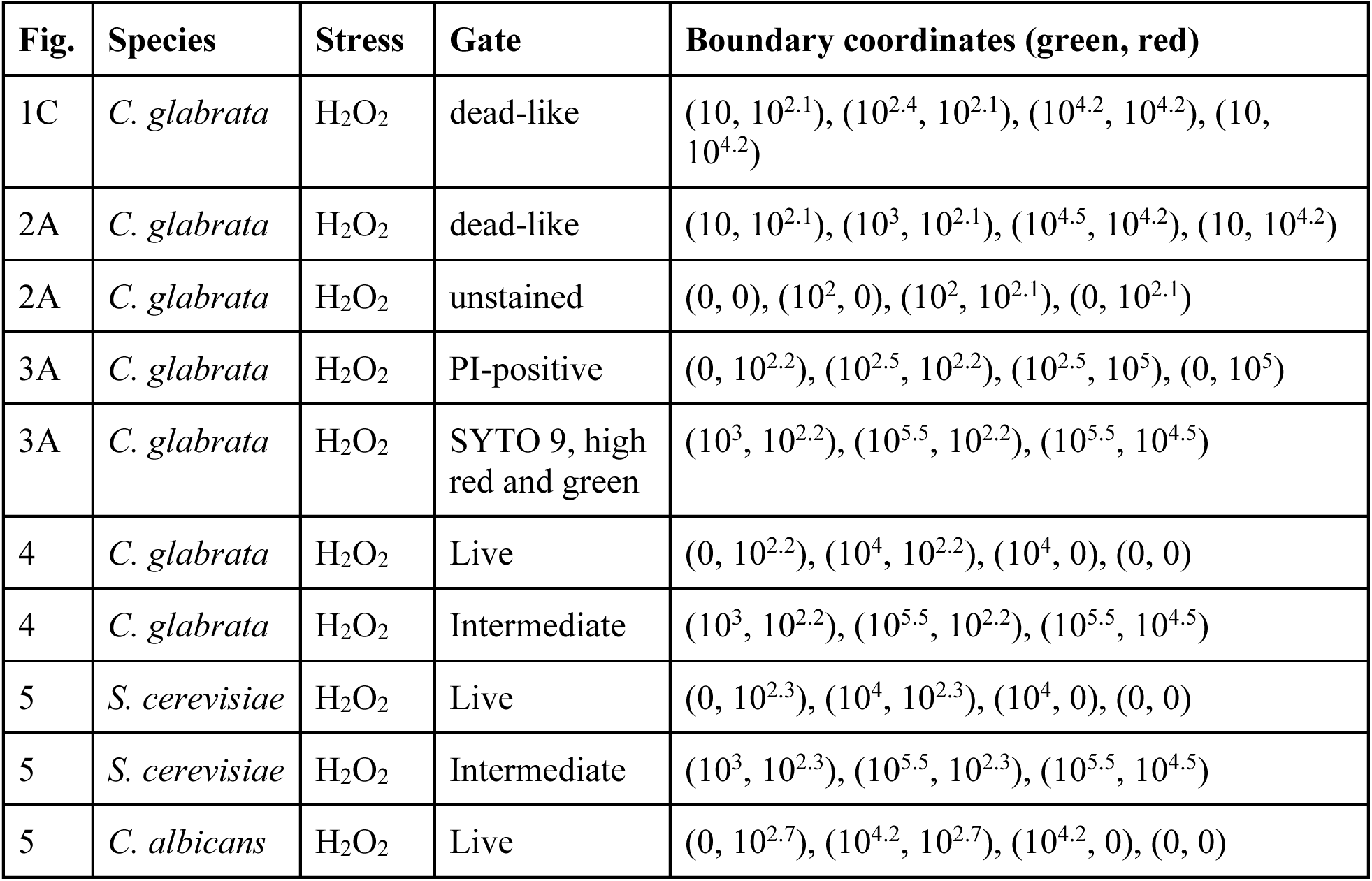
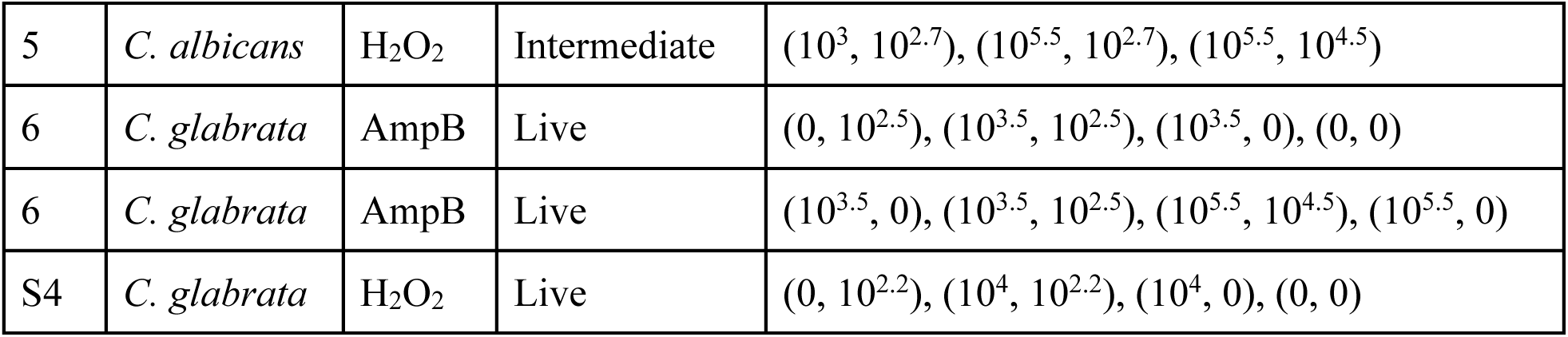
Flow cytometry gate boundaries.

| Fig. | Species | Stress | Gate | Boundary coordinates (green, red) |
| --- | --- | --- | --- | --- |
| 1C | *C. glabrata* | H<sub>2</sub>O<sub>2</sub> | dead-like | (10, 10<sup>2.1</sup>), (10<sup>2.4</sup>, 10<sup>2.1</sup>), (10<sup>4.2</sup>, 10<sup>4.2</sup>), (10, 10<sup>4.2</sup>) |
| 2A | *C. glabrata* | H<sub>2</sub>O<sub>2</sub> | dead-like | (10, 10<sup>2.1</sup>), (10<sup>3</sup>, 10<sup>2.1</sup>), (10<sup>4.5</sup>, 10<sup>4.2</sup>), (10, 10<sup>4.2</sup>) |
| 2A | *C. glabrata* | H<sub>2</sub>O<sub>2</sub> | unstained | (0, 0), (10<sup>2</sup>, 0), (10<sup>2</sup>, 10<sup>2.1</sup>), (0, 10<sup>2.1</sup>) |
| 3A | *C. glabrata* | H<sub>2</sub>O<sub>2</sub> | PI-positive | (0, 10<sup>2.2</sup>), (10<sup>2.5</sup>, 10<sup>2.2</sup>), (10<sup>2.5</sup>, 10<sup>5</sup>), (0, 10<sup>5</sup>) |
| 3A | *C. glabrata* | H<sub>2</sub>O<sub>2</sub> | SYTO 9, high red and green | (10<sup>3</sup>, 10<sup>2.2</sup>), (10<sup>5.5</sup>, 10<sup>2.2</sup>), (10<sup>5.5</sup>, 10<sup>4.5</sup>) |
| 4 | *C. glabrata* | H<sub>2</sub>O<sub>2</sub> | Live | (0, 10<sup>2.2</sup>), (10<sup>4</sup>, 10<sup>2.2</sup>), (10<sup>4</sup>, 0), (0, 0) |
| 4 | *C. glabrata* | H<sub>2</sub>O<sub>2</sub> | Intermediate | (10<sup>3</sup>, 10<sup>2.2</sup>), (10<sup>5.5</sup>, 10<sup>2.2</sup>), (10<sup>5.5</sup>, 10<sup>4.5</sup>) |
| 5 | *S. cerevisiae* | H<sub>2</sub>O<sub>2</sub> | Live | (0, 10<sup>2.3</sup>), (10<sup>4</sup>, 10<sup>2.3</sup>), (10<sup>4</sup>, 0), (0, 0) |
| 5 | *S. cerevisiae* | H<sub>2</sub>O<sub>2</sub> | Intermediate | (10<sup>3</sup>, 10<sup>2.3</sup>), (10<sup>5.5</sup>, 10<sup>2.3</sup>), (10<sup>5.5</sup>, 10<sup>4.5</sup>) |
| 5 | *C. albicans* | H<sub>2</sub>O<sub>2</sub> | Live | (0, 10<sup>2.7</sup>), (10<sup>4.2</sup>, 10<sup>2.7</sup>), (10<sup>4.2</sup>, 0), (0, 0) |
| 5 | *C. albicans* | H<sub>2</sub>O<sub>2</sub> | Intermediate | (10<sup>3</sup>, 10<sup>2.7</sup>), (10<sup>5.5</sup>, 10<sup>2.7</sup>), (10<sup>5.5</sup>, 10<sup>4.5</sup>) |
| 6 | *C. glabrata* | AmpB | Live | (0, 10<sup>2.5</sup>), (10<sup>3.5</sup>, 10<sup>2.5</sup>), (10<sup>3.5</sup>, 0), (0, 0) |
| 6 | *C. glabrata* | AmpB | Live | (10<sup>3.5</sup>, 0), (10<sup>3.5</sup>, 10<sup>2.5</sup>), (10<sup>5.5</sup>, 10<sup>4.5</sup>), (10<sup>5.5</sup>, 0) |
| S4 | *C. glabrata* | H<sub>2</sub>O<sub>2</sub> | Live | (0, 10<sup>2.2</sup>), (10<sup>4</sup>, 10<sup>2.2</sup>), (10<sup>4</sup>, 0), (0, 0) |

### Statistical analyses and data plotting

Figure 3B: Significant differences in the percent SYTO 9-intermediate gate events between the H_2_O_2_-treated samples and the mock were determined by a linear model, i.e., %intermediate ∼ [H_2_O_2_]. H_2_O_2_ doses were treated as a categorical variable, with mock as the reference (intercept). The effect of each of the three non-zero doses was estimated by the linear model. Statistical significance was determined using a Student’s t-test. The two-sided *p*-values were corrected for multiple testing using the Holm-Bonferroni procedure and a test was considered significant when the corrected *p*-value was lower than 0.05. The same procedure was applied to PI-positive events. Figure 4<u>, 5, 6, S4</u>: Pairwise comparisons of either percent live or percent intermediate events were performed using Tukey’s range test. A *p*-value of smaller than 0.05 was considered significant and shown as a filled circle in the figures.

## Acknowledgement

We would like to thank past and current members of the Gene Regulatory Lab for reading and giving feedback on the manuscript. We also want to thank Drs. Jan Fassler and Ana Llopart for reading an early draft of the paper and giving valuable feedback.

## Data Availability

All raw data, including CFU assay and FungaLight staining results, are available at https://github.com/binhe-lab/E041-yeast-viability-assay/releases/tag/preprint. At the time of publication, this repository will be archived with Zenodo to generate a DOI.

## Funding

This work is supported by NIH R35GM137831 and a startup fund from the University of Iowa to BZH. HT received summer support from the University of Iowa Provost Office’s Investment in Strategic Priorities fund. JL received summer support from the Carol B. and Robert G. Lynch Department of Biology Graduate Fund at the University of Iowa.

## Supporting Information

**Fig. S1.**
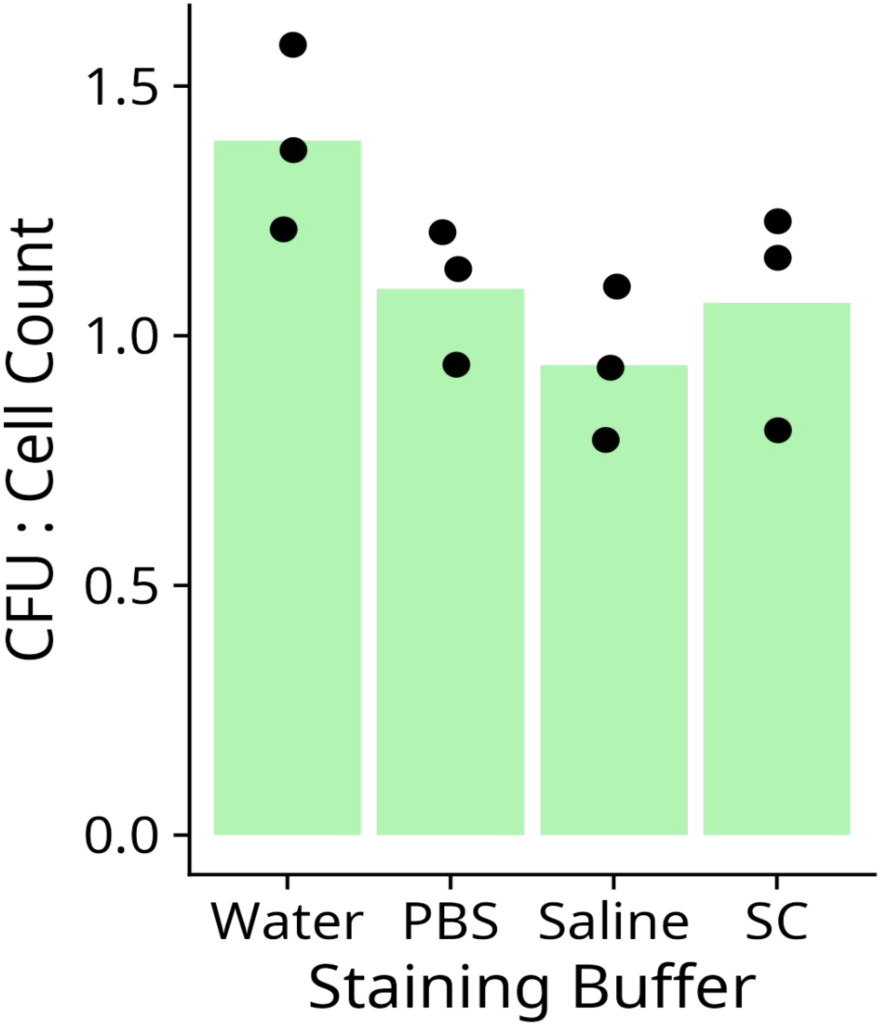
Viability of mock-treated mid-log phase *C. glabrata* cells. Ratios between the CFU and the number of cells plated based on flow cytometry in mock-treated (mid-log) *C. glabrata* cells resuspended in the indicated buffers. Dots represent the individual biological replicate and bars represent the mean.

**Fig. S2.**
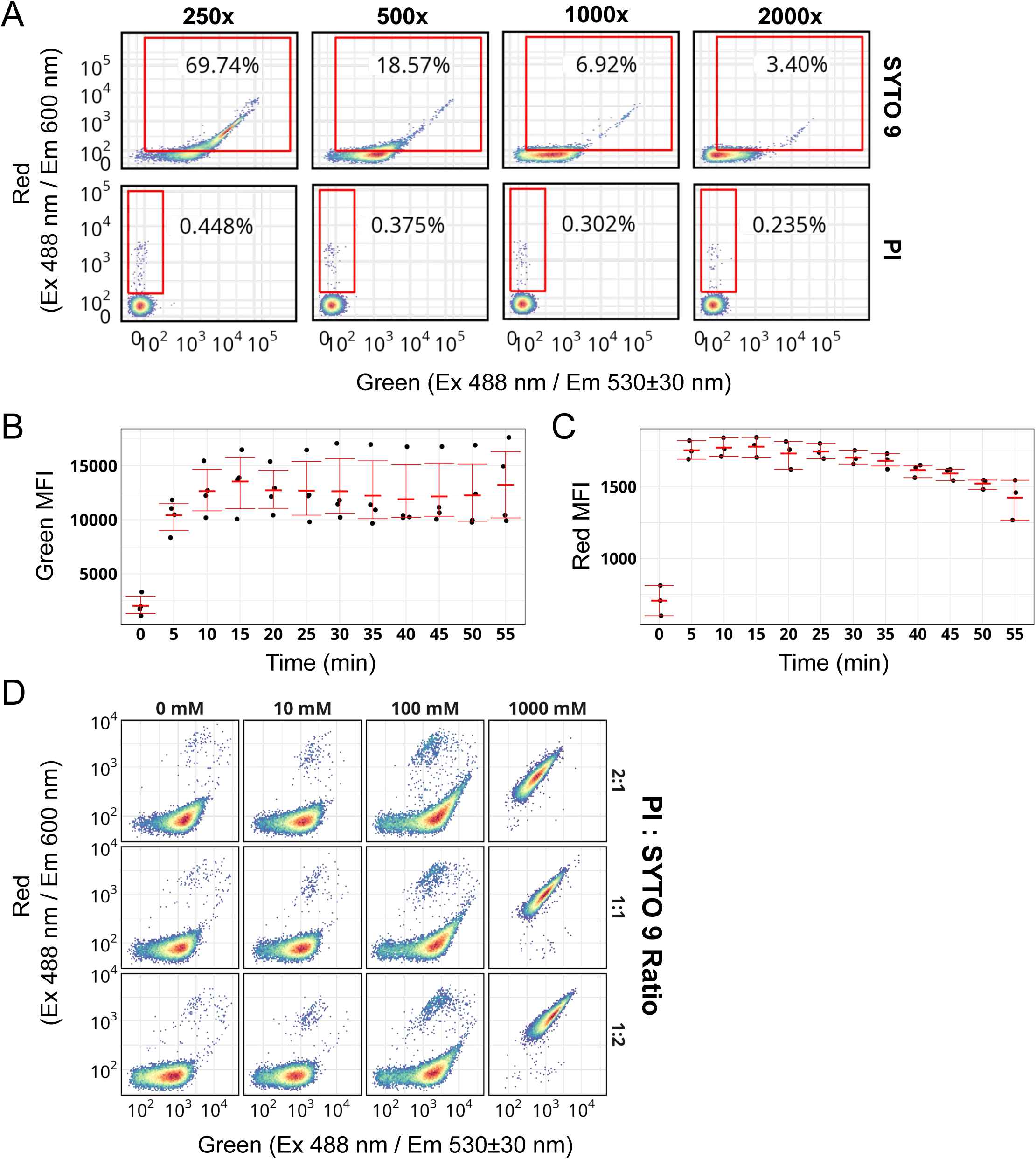
Testing dye concentrations, incubation time and dye ratio on the staining pattern. (A) 2D density plots showing staining patterns of mid-log *C. glabrata* cells at various dilutions of SYTO 9 or PI (stock concentrations: 3.34 mM for SYTO 9, 20 mM for PI). Rectangular gates were used to quantify cell events considered damaged or dead. (B) Median Fluorescence Intensity (MFI) over time for SYTO 9 stained mid-log *C. glabrata* cells. Dots show individual biological replicates (n = 4, biological replicates). Thick red bars represent the mean and the error bars represent the 95% confidence intervals. (C) Similar to B for PI-stained, heat-killed *C. glabrata* cells. (D) 2D density plots for *C. glabrata* cells treated with H_2_O_2_ and stained with various ratios of PI:SYTO 9 (right). For all samples, SYTO 9 is applied at a fixed concentration of 3.34 μM; PI is used at 20 μM at 1:1 ratio.

**Fig. S3.**
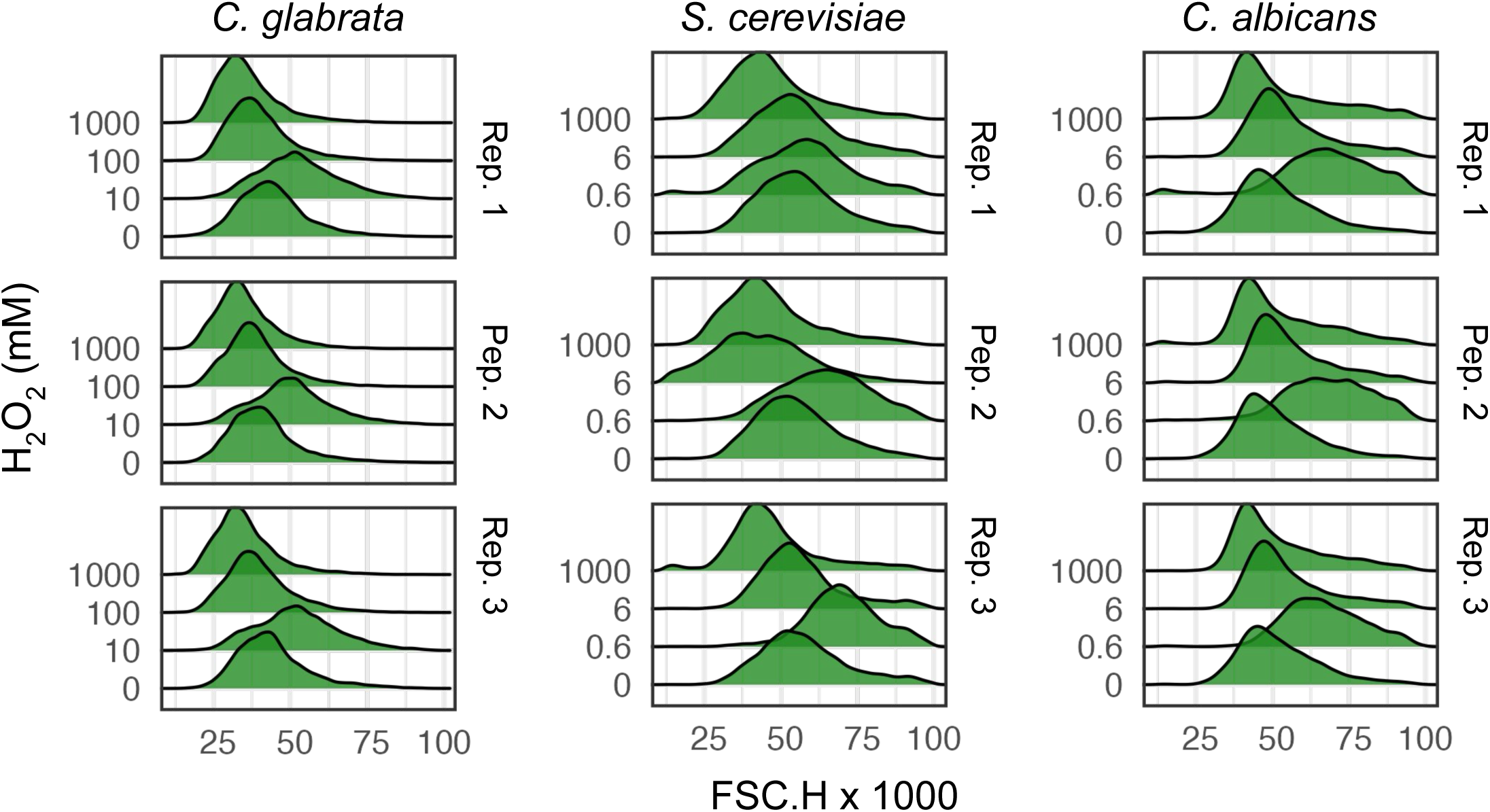
H_2_O_2_ treatment causes changes in cell size as indicated by FSC (forward scattering). FSC.H (height) distribution of gated single cell events were shown for each of the three species as indicated on the top. Cells were treated for 2 hours at four different H_2_O_2_ doses (y-axis labels). Three biological replicates were shown for each species and treatment.

**Fig. S4.**
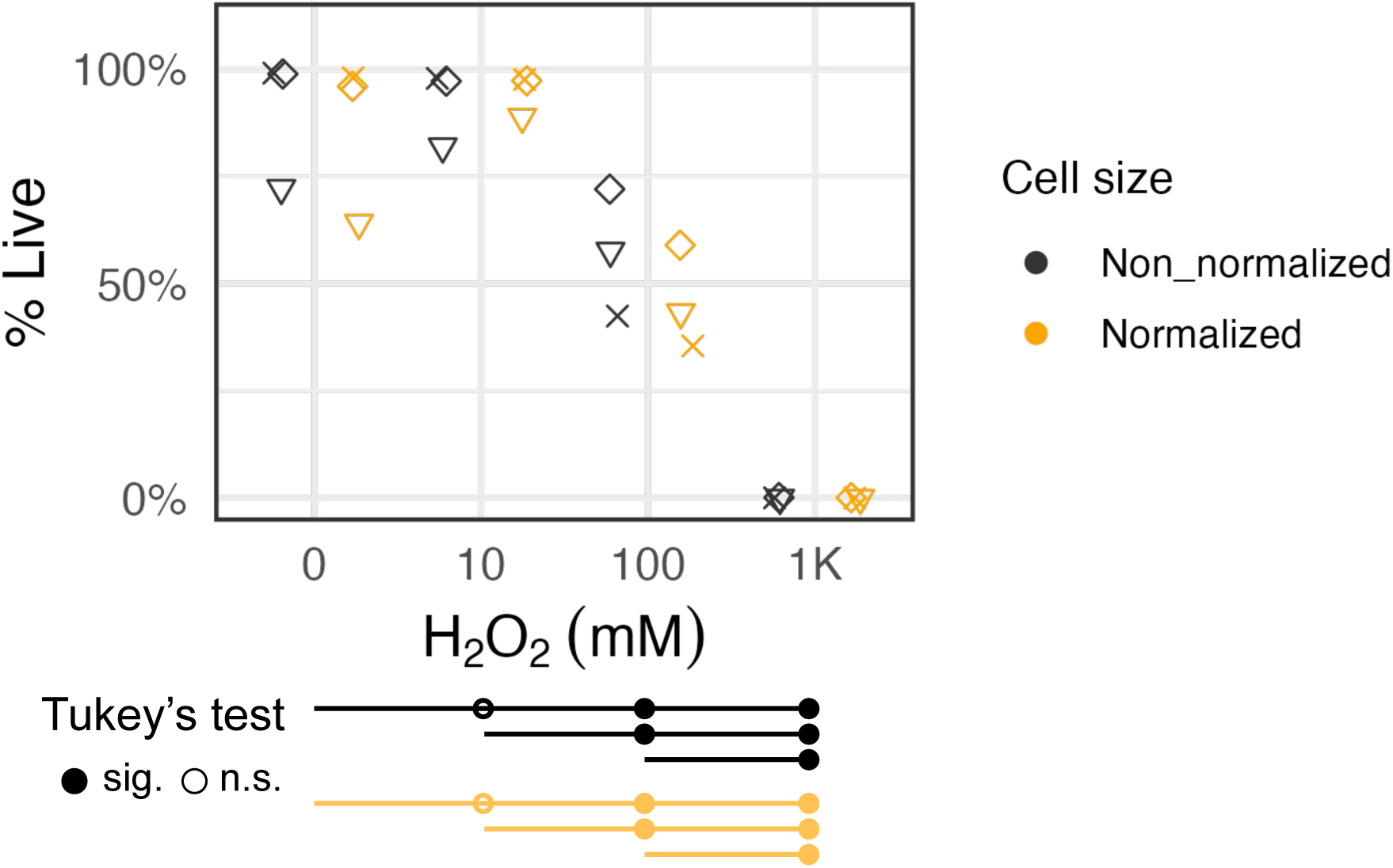
Normalizing fluorescence by FSC.H didn’t reduce the sample variance dramatically.. Fluorescence signals were first normalized by FSC.H (see Methods for details) and then the same gating strategies were applied as for the unnormalized data to calculate the percent live values. The resulting estimates were plotted side-by-side with the non-normalized (Fig. 4B) results for *C. glabrata* treated with different concentrations of H_2_O_2_. Statistical test results were shown below the graph similarly to Fig. 4B.

**Fig. S5.**
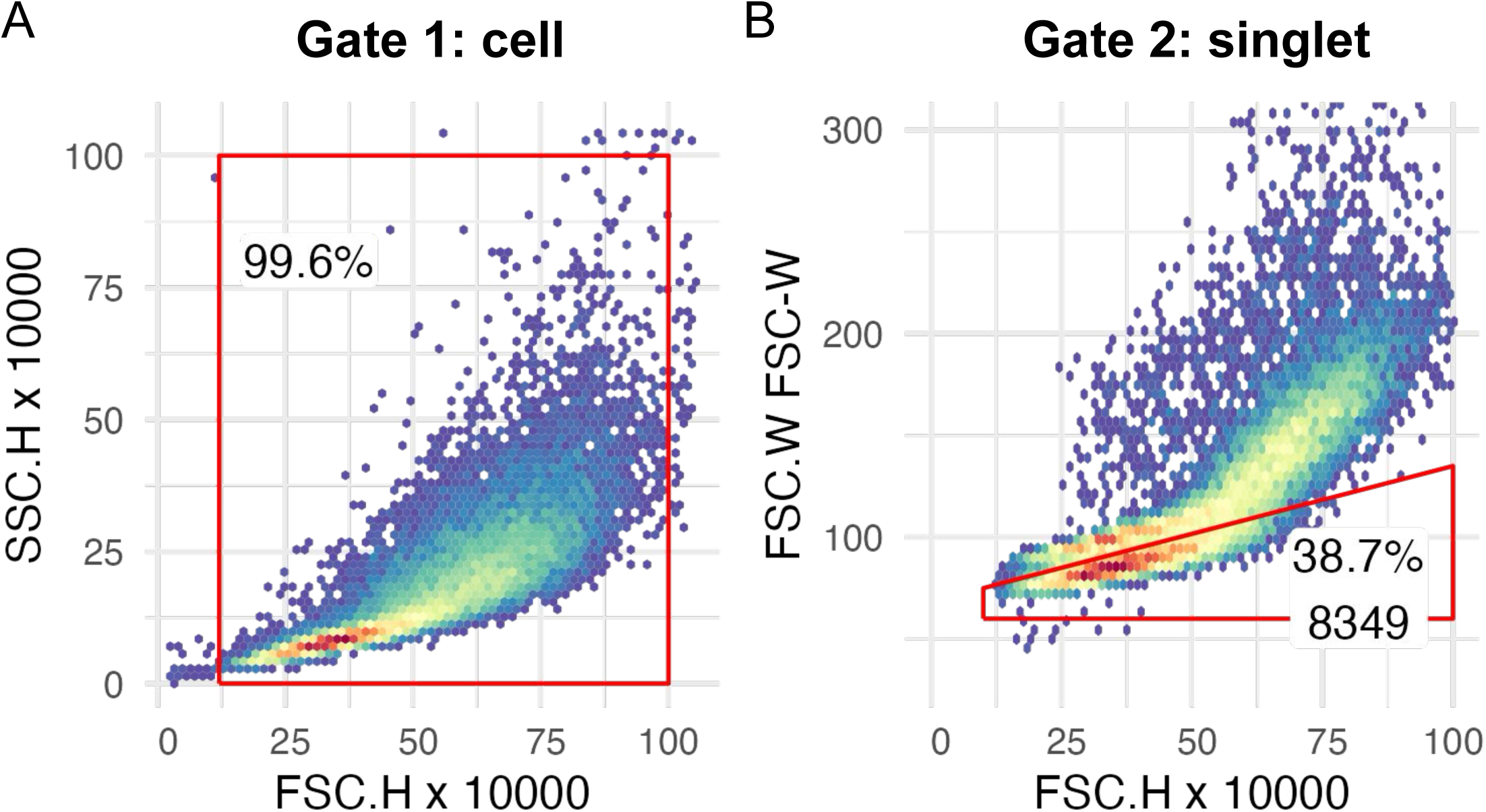
Example gating strategy for flow cytometry data. (A) All events were plotted on FSC.H and SSC.H. A rectangular gate is used to exclude non-cell events. (B) Events within the first gate (“cell”) were plotted on FSC.H and FSC.W. Singlets (single cell event as opposed to doublets or multilets) were selected by excluding events with a higher FSC.W given the same FSC.H.

## Notes

https://github.com/binhe-lab/E041-yeast-viability-assay

### Competing Interest Statement

The authors have declared no competing interest.

### Summary of Updates

revised text and enhanced the introduction.

## Literature Cited

1. Lopes JP, Lionakis MS. Pathogenesis and virulence of Candida albicans. Virulence. 13: 89–121. doi:10.1080/21505594.2021.2019950

2. Hazen KC. New and emerging yeast pathogens. Clin Microbiol Rev. 1995;8: 462–478.

3. Welsh RM, Bentz ML, Shams A, Houston H, Lyons A, Rose LJ, et al. Survival, Persistence, and Isolation of the Emerging Multidrug-Resistant Pathogenic Yeast Candida auris on a Plastic Health Care Surface. J Clin Microbiol. 2017;55: 2996–3005. doi:10.1128/jcm.00921-17

4. Quinto-Alemany D, Canerina-Amaro A, Hernández-Abad LG, Machín F, Romesberg FE, Gil-Lamaignere C. Yeasts Acquire Resistance Secondary to Antifungal Drug Treatment by Adaptive Mutagenesis. PLoS ONE. 2012;7: e42279. doi:10.1371/journal.pone.0042279

5 . Novak Babič M, Zalar P, Ženko B, Džeroski S, Gunde-Cimerman N. Yeasts and yeast-like fungi in tap water and groundwater, and their transmission to household appliances. Fungal Ecol. 2016;20: 30–39. doi:10.1016/j.funeco.2015.10.001

6. Lourens-Hattingh A, Viljoen BC. Growth and survival of a probiotic yeast in dairy products. Food Res Int. 2001;34: 791–796. doi:10.1016/S0963-9969(01)00085-0

7. Blagosklonny MV. Cell senescence: Hypertrophic arrest beyond the restriction point. J Cell Physiol. 2006;209: 592–597. doi:10.1002/jcp.20750

8. Davey HM. Life, Death, and In-Between: Meanings and Methods in Microbiology. Appl Environ Microbiol. 2011;77: 5571–5576. doi:10.1128/AEM.00744-11

9. Carmona-Gutierrez D, Bauer MA, Zimmermann A, Aguilera A, Austriaco N, Ayscough K, et al. Guidelines and recommendations on yeast cell death nomenclature. Microb Cell. 2018;5: 4–31. doi:10.15698/mic2018.01.607

10. Wloch-Salamon DM, Bem AE. Types of cell death and methods of their detection in yeast Saccharomyces cerevisiae. J Appl Microbiol. 2013;114: 287–298. doi:10.1111/jam.12024

11. Kwolek-Mirek M, Zadrag-Tecza R. Comparison of methods used for assessing the viability and vitality of yeast cells. FEMS Yeast Res. 2014;14: 1068–1079. doi:10.1111/1567-1364.12202

12. Sahu SR, Utkalaja BG, Patel SK, Acharya N. Spot Assay and Colony Forming Unit (CFU) Analyses-based sensitivity test for Candida albicans and Saccharomyces cerevisiae. Bio-Protoc. 2023;13: e4872. doi:10.21769/BioProtoc.4872

13. Olivares-Marin IK, González-Hernández JC, Regalado-Gonzalez C, Madrigal-Perez LA. Saccharomyces cerevisiae Exponential Growth Kinetics in Batch Culture to Analyze Respiratory and Fermentative Metabolism. J Vis Exp JoVE. 2018; 58192. doi:10.3791/58192

14. Khan A ul M, Torelli A, Wolf I, Gretz N. AutoCellSeg: robust automatic colony forming unit (CFU)/cell analysis using adaptive image segmentation and easy-to-use post-editing techniques. Sci Rep. 2018;8: 7302. doi:10.1038/s41598-018-24916-9

15. Boukouvalas DT, Prates RA, Lima Leal CR, de Araújo SA. Automatic segmentation method for CFU counting in single plate-serial dilution. Chemom Intell Lab Syst. 2019;195: 103889. doi:10.1016/j.chemolab.2019.103889

16. Stolze N, Bader C, Henning C, Mastin J, Holmes AE, Sutlief AL. Automated image analysis with ImageJ of yeast colony forming units from cannabis flowers. J Microbiol Methods. 2019;164: 105681. doi:10.1016/j.mimet.2019.105681

17. Kwolek-Mirek M, Zadrag-Tecza R. Comparison of methods used for assessing the viability and vitality of yeast cells. FEMS Yeast Res. 2014;14: 1068–1079. doi:10.1111/15671364.12202

18. Sun S, Baryshnikova A, Brandt N, Gresham D. Genetic interaction profiles of regulatory kinases differ between environmental conditions and cellular states. Mol Syst Biol. 2020;16: e9167. doi:10.15252/msb.20199167

19. Zhang T, Fang HHP. Quantification of Saccharomyces cerevisiae viability using BacLight. Biotechnol Lett. 2004;26: 989–992. doi:10.1023/b:bile.0000030045.16713.19

20. Stocks SM. Mechanism and use of the commercially available viability stain, BacLight. Cytometry A. 2004;61A: 189–195. doi:10.1002/cyto.a.20069

21. Boulos L, Prévost M, Barbeau B, Coallier J, Desjardins R. LIVE/DEAD BacLight : application of a new rapid staining method for direct enumeration of viable and total bacteria in drinking water. J Microbiol Methods. 1999;37: 77–86. doi:10.1016/s0167-7012(99)00048-2

22. Jin Y, Zhang T, Samaranayake YH, Fang HHP, Yip HK, Samaranayake LP. The use of new probes and stains for improved assessment of cell viability and extracellular polymeric substances in Candida albicans biofilms. Mycopathologia. 2005;159: 353–360. doi:10.1007/s11046-004-6987-7

23. Bojsen R, Regenberg B, Folkesson A. Saccharomyces cerevisiae biofilm tolerance towards systemic antifungals depends on growth phase. BMC Microbiol. 2014;14: 305. doi:10.1186/s12866-014-0305-4

24. Ryan LK, Freeman KB, Masso-Silva JA, Falkovsky K, Aloyouny A, Markowitz K, et al. Activity of potent and selective host defense peptide mimetics in mouse models of oral candidiasis. Antimicrob Agents Chemother. 2014;58: 3820–3827. doi:10.1128/AAC.02649-13

25. Roscetto E, Contursi P, Vollaro A, Fusco S, Notomista E, Catania MR. Antifungal and anti-biofilm activity of the first cryptic antimicrobial peptide from an archaeal protein against Candida spp. clinical isolates. Sci Rep. 2018;8: 17570. doi:10.1038/s41598-018-35530-0

26. Chen Y-C, Yang Y, Zhang C, Chen H-Y, Chen F, Wang K-J. A Novel Antimicrobial Peptide Sparamosin26-54 From the Mud Crab Scylla paramamosain Showing Potent Antifungal Activity Against Cryptococcus neoformans. Front Microbiol. 2021;12: 746006. doi:10.3389/fmicb.2021.746006

27. Wainschtein P, Jain D, Zheng Z, TOPMed Anthropometry Working Group, NHLBI Trans-Omics for Precision Medicine (TOPMed) Consortium, Cupples LA, et al. Assessing the contribution of rare variants to complex trait heritability from whole-genome sequence data. Nat Genet. 2022;54: 263–273. doi:10.1038/s41588-021-00997-7

28. Chudzik B, Koselski M, Czuryło A, Trębacz K, Gagoś M. A new look at the antibiotic amphotericin B effect on Candida albicans plasma membrane permeability and cell viability functions. Eur Biophys J EBJ. 2015;44: 77–90. doi:10.1007/s00249-014-1003-8

29. Sun J, Liu X, Jiang G, Qi Q. Inhibition of Nucleic Acid Biosynthesis Makes Little Difference to Formation of Amphotericin B-Tolerant Persisters in Candida albicans Biofilm. Antimicrob Agents Chemother. 2015;59: 1627–1633. doi:10.1128/aac.03765-14

30. Ponomarova O, Gabrielli N, Sévin DC, Mülleder M, Zirngibl K, Bulyha K, et al. Yeast Creates a Niche for Symbiotic Lactic Acid Bacteria through Nitrogen Overflow. Cell Syst. 2017;5: 345–357.e6. doi:10.1016/j.cels.2017.09.002

31. Stiefel P, Schmidt-Emrich S, Maniura-Weber K, Ren Q. Critical aspects of using bacterial cell viability assays with the fluorophores SYTO9 and propidium iodide. BMC Microbiol. 2015;15: 36. doi:10.1186/s12866-015-0376-x

32. Rego A, Ribeiro A, Côrte-Real M, Chaves SR. Monitoring yeast regulated cell death: trespassing the point of no return to loss of plasma membrane integrity. Apoptosis. 2022;27: 778–786. doi:10.1007/s10495-022-01748-7

33. Nikolaou E, Agrafioti I, Stumpf M, Quinn J, Stansfield I, Brown AJ. Phylogenetic diversity of stress signalling pathways in fungi. BMC Evol Biol. 2009;9: 44. doi:10.1186/1471-2148-9-44

34. Pappas PG, Lionakis MS, Arendrup MC, Ostrosky-Zeichner L, Kullberg BJ. Invasive candidiasis. Nat Rev Dis Primer. 2018;4: 18026. doi:10.1038/nrdp.2018.26

35. Giannuzzi L, Lombardo T, Juárez I, Aguilera A, Blanco G. A Stochastic Characterization of Hydrogen Peroxide-Induced Regulated Cell Death in Microcystis aeruginosa. Front Microbiol. 2021;12: 636157. doi:10.3389/fmicb.2021.636157

36. Cormack BP, Falkow S. Efficient homologous and illegitimate recombination in the opportunistic yeast pathogen Candida glabrata. Genetics. 1999;151: 979–987.

37. Homann OR, Dea J, Noble SM, Johnson AD. A Phenotypic Profile of the Candida albicans Regulatory Network. PLOS Genet. 2009;5: e1000783. doi:10.1371/journal.pgen.1000783

